# Extracellular matrix and cyclic stretch alter fetal cardiomyocyte proliferation and maturation in a rodent model of heart hypoplasia

**DOI:** 10.1101/2020.08.17.254334

**Authors:** Matthew C. Watson, Corin Williams, Raymond M. Wang, Luke R. Perreault, Kelly E. Sullivan, Whitney L. Stoppel, Lauren D. Black

## Abstract

Birth defects, particularly those that affect development of the heart, are a leading cause of morbidity and mortality in infants and young children. Babies born with heart hypoplasia (HH) disorders often have a poor prognosis. It remains unclear whether cardiomyocytes from hypoplastic hearts retain the potential to recover growth, although this knowledge would be beneficial for developing therapies for HH disorders. The objective of this study was to determine the proliferation and maturation potential of cardiomyocytes from hypoplastic hearts and whether these behaviors are influenced by biophysical signaling from the extracellular matrix (ECM) and cyclic mechanical stretch. Congenital diaphragmatic hernia (CDH)-associated HH was induced in rat fetuses by maternal exposure to nitrofen. Hearts were isolated from embryonic day 21 nitrofen-treated fetuses positive for CDH (CDH+) and from fetuses without nitrofen administration during gestation. CDH+ hearts were smaller and had decreased myocardial proliferation, along with evidence of decreased maturity compared to healthy hearts. In culture, CDH+ cardiomyocytes remained immature and demonstrated increased proliferative capacity compared to their healthy counterparts. Culture on ECM derived from CDH+ hearts led to a significant reduction in proliferation for both CDH+ and healthy cardiomyocytes. Healthy cardiomyocytes were dosed with exogenous nitrofen to examine whether nitrofen may have an abhorrent effect on the proliferative ability of cardiomyocyte, yet no significant change in proliferation was observed. When subjected to stretch, CDH+ cardiomyocytes underwent lengthening of sarcomeres while healthy cardiomyocyte sarcomeres were unaffected. Taken together, our results suggest that alterations to environmental cues such as ECM and stretch may be important factors in the pathological progression of HH.

## 1. Introduction

Congenital diaphragmatic hernia (CDH) is a serious birth defect that occurs in approximately 1 in 2500-3000 live births. Only ∼16% of prenatally diagnosed CDH patients are expected to survive past the first year of life [1]. Failure of the diaphragm to close permits intrusion of visceral organs into the thoracic cavity and subsequent compression of the developing heart and lungs. Heart and/or lung hypoplasia is associated with a particularly poor prognosis in CDH [2]. Interestingly, patients born with mild to moderate heart hypoplasia (HH) exhibit restored heart growth after surgical repair of CDH [3]; however, the underlying mechanisms are unknown. A better understanding of the factors that influence heart growth during development is not only critical for improving therapies for children with heart defects, but would be invaluable to the field of cardiac regeneration as a whole.

Cardiomyocyte proliferation plays a key role in cardiac growth during fetal development [4] and may provide an explanation for restored heart growth in repaired CDH. In other forms of HH, such as Hypoplastic Left Heart Syndrome (HLHS), there is evidence that cardiomyocyte proliferation is severely diminished [5], although this has not been established in CDH-associated HH. Cardiomyocyte behavior can be regulated by a variety of biophysical cues, such as the extracellular matrix (ECM) [6, 7], growth factors [6, 8], cell-cell contacts [9], substrate stiffness [10, 11], and mechanical stretch [12]. Although current data is limited, studies suggest that at least some of these signals may be altered in the developing hypoplastic heart [21].

It is intriguing to consider the possibility that cardiomyocyte proliferation can be recovered in HH if pathological conditions are removed or healthy biophysical environments are restored. The experimental manipulation of mechanical loading in embryonic zebrafish [13] and chick hearts [14] motivated fetal aortic annuloplasty in severe HLHS in an attempt to restore normal blood flow and, subsequently, left heart growth [15]. We hypothesized that cardiomyocyte proliferation is reduced in CDH-associated HH, but that the cells retain the ability to proliferate if they are removed from the hypoplastic environment. We used the nitrofen model of CDH in rats, which is thought to be similar in etiology and phenotype to human CDH [16-18]. In addition to studying proliferation in native hearts, we isolated cardiac cells and assessed their response to two external cues that have been implicated in HH defects: ECM, which is thought to be structurally immature in HH [19, 20], and cyclic stretch, which is diminished in HH [12]. Our *in vitro* culture systems allowed us to systematically study these cues independently, which would not be possible *in vivo*. We found that CDH+ cardiomyocytes were more proliferative than healthy cardiomyocytes when placed in culture, and that ECM and cyclic stretch (1 Hz, 5% amplitude) differentially regulated proliferation and maturation.

## 2. Materials and Methods

### 2.1. Nitrofen model of congenital diaphragmatic hernia

All animal procedures were performed in accordance with the Institutional Animal Care and Use Committee at Tufts University and the NIH Guide for the Care and Use of Laboratory Animals. Pregnant Sprague-Dawley rats at gestational day E10 (purchased from Charles River Laboratories, Wilmington, MA) were subjected to short Isoflurane anesthesia and immediately gavaged with a single dose of 100 mg nitrofen (Wako Pure Chemical Industries, Japan) dissolved in 2 ml olive oil. Control animals received olive oil alone, similar to previously described methods [21]. Pregnancy then progressed until E21, at which point the dams were sacrificed by CO_2_ inhalation. The fetuses were harvested for the studies described below.

### 2.2. Fetal heart harvest

Immediately after harvest, fetuses were placed on ice, and euthanized by decapitation. The chest wall was carefully opened above the diaphragm and the heart and lungs were removed. The diaphragm was then checked for the presence of CDH. Nitrofen treatment at E10 resulted in CDH in approximately 80-85% of the fetuses, similar to what others have found [22]. As so few nitrofen-treated CDH negative hearts were found (approximately 2-4 out of 12-16 hearts per litter), they were not included in the present study. CDH was never detected in control fetuses. Nitrofen-treated CDH positive (CDH+) and control (“healthy”) hearts were weighed prior to further characterization.

### 2.3. Histology and imaging of native heart sections

Whole hearts were fixed by immersion in 4% paraformaldehyde at 4°C overnight. After washing with phosphate buffered saline (PBS), the samples were cryo-protected in sucrose solution (30% wt/vol), and then embedded in Tissue Tek optimum cutting temperature (OCT) compound. The hearts were sectioned (thickness of 7 µm) on a Leica CM 1950 CryoStat. Heart slices were stored at −20°C until use. OCT compound was removed by washing with PBS prior to staining with hematoxylin and eosin (H&E) to visualize gross heart structure. Hearts were imaged on a Keyence BZ-X700 fluorescent microscope using a color camera. The thickness of the compact zone of the left ventricular free wall was measured with ImageJ using H&E stained sections.

### 2.4. ECM composition

A subset of CDH+ and healthy hearts were used to determine extracellular matrix (ECM) composition by liquid chromatography tandem mass spectrometry (LC-MS/MS), similar to our previous studies of cardiac ECM [7]. Immediately after weighing, freshly isolated whole hearts were decellularized in 0.1% sodium dodecyl sulfate (SDS) for 48 hr at room temperature with agitation on an orbital shaker. The resulting ECM was then incubated in 0.1% TritonX-100 (Amresco, Solon, OH) for ∼3-4 hr, washed with distilled water, frozen, and lyophilized overnight (Labconco, Kansas City, MO). The ECM of individual fetal hearts was solubilized in 200 μl urea solution (5 M urea, 2 M thiourea, 50 mM DTT, 0.1% SDS) [23] for 20 hr with agitation. Finally, the ECM was precipitated in acetone and analyzed via LC-MS/MS within 24 hr at the Beth Israel Deaconess Medical Center Mass Spectrometry Core Facility. Among the samples (N = 3 for each condition), 34 unique ECM components were identified and their relative percentages in the total ECM composition was determined from spectrum counts.

### 2.5. Immunocytochemistry

To study proliferation and sarcomere development in the native healthy and CDH+ hearts, sections were stained for nuclei (Hoechst 33258), Ki67 (Abcam, rabbit polyclonal), and sarcomeric α-actinin (Sigma, mouse monoclonal). Briefly, the samples were washed with PBS, then blocked with 5% donkey serum and 1% bovine serum albumin (BSA) for 1 hr at room temperature. Incubation with primary antibodies was for 1 hr, followed by 3 PBS washes, then incubation with secondary antibodies (AlexaFluor 488 donkey anti-rabbit IgG, AlexaFluor 555 donkey anti-mouse IgG; Invitrogen) for 1 hr. After washing with PBS, the samples were imaged on an Olympus IX8I microscope with Metamorph Basic software (version 7.7.4.0, Molecular Devices). Image analysis to determine cardiomyocyte proliferation and sarcomere length was carried out as described below in sections 2.8 and 2.9.

### 2.6. Cardiac gene expression by quantitative PCR

A panel of cardiac genes was analyzed by PCR. RNA was isolated from fetal hearts using the RNeasy® Mini Kit (Qiagen Sciences, Germantown, MD) per the manufacturer’s instructions. RNA concentration was determined using a Nanodrop 2000 Spectrophotometer (Thermo Scientific, Waltham, MA) and purity was assessed by the ratio of the 260/280 absorbance readings. Samples with high purity were then used to make cDNA using the High Capacity cDNA Reverse Transcription Kit (Applied Biosystems, Foster City, CA) per the manufacturer’s instructions. cDNA was loaded into the wells of a MicroAmp® Optical 96-well reaction plate with 10 µL of 2X TaqMan® Gene Expression Master Mix and 1 µL of predesigned 20X TaqMan® Gene Expression Assay primers for the specific gene of interest (Applied Biosystems, Foster City, CA) diluted in nuclease-free water to a final volume of 20 µL. Samples were probed for gene expression related to contractile function (MYH6, MYH7, TNNT2, TNNI3, ACTN2, ATPA2), gap junction signaling (GJA1, CDH2) and early markers of the cardiac lineage (GATA4, GATA6, NKX2-5). Expression of each gene was normalized to GAPDH values. Quantitative RT-PCR was performed using the Mx3000P QPCR System (Agilent Technologies, Lexington, MA) with incubation parameters of 2 min at 50°C, 10 min at 95°C and 40 cycles of 15 seconds at 95°C followed by 1 min at 60°C. Ct values were determined by the software provided with the Mx3000P QPCR System and differences in mRNA expression were calculated by the 2^-ΔΔct^ method [24] based on validation tests performed by Applied Biosystems.

### 2.7. Cardiac cell isolation and culture

Cells from CDH+ and healthy control hearts were isolated according to our previously described methods [25]. Briefly, hearts were isolated from euthanized fetal pups at E21, the ventricles were minced, and the tissue underwent 7 × 7 min digestions in collagenase type II (Worthington Biochemical Corp, Lakewood, NJ) and sterile PBS supplemented with 20 mM glucose. Cells were counted with a hemocytometer and seeded at a density of 100,000 cells/cm^2^ into tissue culture polystyrene 48-well plates. The culture medium contained 15% fetal bovine serum (FBS) in Dulbecco’s Modified Eagle Medium (DMEM) with 1% penicillin-streptomycin and was changed every 2 days. The cells were fixed and stained with Hoechst, Ki67, and α-actinin at various time points, imaged, and analyzed as described below in sections 2.8 and 2.9.

### 2.8. Cell proliferation measurements

Cell numbers and proliferation were determined using custom pipelines in CellProfiler (release 11710, the Broad Institute). Total cell nuclei were determined from the Hoechst stain and proliferating cells (Ki67+ nuclei) were determined from the Ki67 stain. The α-actinin stain used to label cardiomyocytes was used as a “mask” to identify cardiomyocyte-specific nuclei and proliferation. Total cell and cardiomyocyte density was calculated using the total imaged area (converted to mm^2^) for each sample. Cardiomyocyte-specific proliferation was measured as the percentage of proliferating cells that were also positive for sarcomeric α-actinin ((Ki67+ α-actinin+)/Ki67+).

### 2.9. Cardiac cell culture with exogenous nitrofen

To determine whether nitrofen would affect the behavior of healthy cardiomyocytes, cardiomyocytes freshly isolated from neonatal hearts were seeded onto 12 well plates at a density of 50,000 cells/cm^2^. Cells were cultured in serum containing medium until beating was observed. Once beating was observed, cells were treated with medium containing nitrofen at 50 ug/mL, and 100 ug/mL. Dosages were determined by estimating the concentrations of nitrofen delivered to each rat fetus. Serum containing medium was used as a control. For normalization purposes, a subset of each group was fixed at Cells were treated on days 1 and 2, and wells from each group were fixed on either 1 day prior to the first nitrofen treatment or 3 days after the beginning of nitrofen treatment. Cells were stained, imaged, and analyzed for proliferation as described above in section 2.8.

### 2.10. Sarcomere measurements

Sarcomere length has been used as a measure of cardiomyocyte maturation [10, 26, 27]. To adequately visualize sarcomeres, images of α-actinin staining at 40x magnification were acquired. Analysis was performed using ImageJ software (NIH, version 1.45s). When organized sarcomeres were observed, a line was manually drawn across multiple sarcomeres perpendicular to alignment. The “Plot Profile” function was used to display the staining intensity across the line. The number of sarcomeres in a given length was counted and the average measured sarcomere length was calculated. Sarcomeres were also categorized according to the following definitions: “developing” (sarcomeres measuring <1.8 μm in length); and “mature” (≥ 1.8 μm) [28].

### 2.11 Cardiac cell culture on ECM

ECM from healthy and CDH+ hearts was obtained as described above and solubilized at a concentration of 10 mg/ml in a solution containing 1 mg/ml pepsin and 0.1 M HCl [7, 29]. The ECM solution was neutralized with NaOH, immediately coated onto 48-well plates at a density of 50 µg/cm^2^ and allowed to dry in a sterile tissue culture cabinet overnight. Prior to cell seeding, the ECM was washed 3 times with sterile PBS. Cells freshly isolated from healthy and CDH+ hearts were then seeded onto the coated plates at a density of 100,000 cells/cm^2^. To avoid the potential confounding effects of serum on proliferation, cells were cultured in a serum-free medium that contained the following: 50/50 mixture of DMEM and Ham’s F12 Nutrient Mix (Invitrogen), 0.2% (wt/vol) bovine serum albumin (BSA) (Sigma), 0.5% (vol/vol) insulin–transferrin–selenium-X (Invitrogen) and 1% penicillin–streptomycin (Invitrogen), with 10 mM L -ascorbic acid (Sigma) added fresh at every feeding [30]. Cells were fed on days 1 and 3, and wells from each group were fixed on either day 1 or day 4 in culture. Cells were stained, imaged, and analyzed for proliferation as described above in section 2.8.

### 2.12 Cardiac cell culture with cyclic mechanical stretch

To determine the effects of cyclic mechanical stretch on cardiomyocyte behavior, cells were cultured on a custom-built cell culture membrane stretching device [31]. Fetal cardiac cells isolated at E21 from healthy and CDH+ hearts were seeded at 1 × 10^6^ cells per well in 6-well Bioflex® culture plates. The experimental set-up was similar to previously described methods with slight modifications [31]. The Bioflex® culture plate membranes were pre-coated with collagen type I by the manufacturer. We found that an additional surface treatment of 4 µg/cm^2^ human plasma Fibronectin (Millipore, Billerica MA) in DMEM applied overnight at 37°C with mild orbital plate agitation was necessary to create a consistent surface coating for cell adhesion. Cells were initially cultured under static conditions for 4 days post-isolation to ensure strong adhesion to the membrane and then stretched for 3 days on the custom-built system. Membranes were deformed based on a standard left ventricular volumetric loading waveform with 1 Hz frequency and 5% amplitude. Control samples were not subjected to stretching (“static”). The underside of the Bioflex® culture membranes was lubricated with a silicone lubricant (Loctite, Düsseldorf, Germany) to minimize friction with the plunger during stretch. Cells were fed culture medium containing 10% horse serum, 2% FBS, and 1% penicillin-streptomycin in DMEM. Medium was changed and lubricant was reapplied every two days. At the end of the experiment, samples were fixed, stained, and imaged as described above. Analysis included proliferation and sarcomere measurements as described above in sections 2.8 and 2.9.

### 2.13 Statistical Analysis

Statistical analysis was performed using analysis of variance (ANOVA) and Tukey’s post-hoc test or the Student’s t-test, as appropriate. Differences were considered statistically significant for p < 0.05.

## 3. Results

### 3.1. Altered gross morphology in CDH+ hearts

Upon isolation at E21, we found that CDH+ hearts were smaller and had significantly decreased mass compared to healthy controls (*Figure 1A, B*). Qualitatively, H & E staining suggested that the CDH+ hearts were morphologically immature, with many still exhibiting the ventricular groove, thinner/ less compacted ventricular walls, and more trabeculation compared to healthy hearts (*Figure 1C*). Measurements of the compact zone of the left ventricular free wall indicated that CDH+ hearts had thinner compact myocardium compared to healthy hearts (400 ± 96 μm vs. 580 ± 48 μm; *Figure 1D, p< 0.01*). ECM was also altered in CDH+ hearts as determined by LC-MS/MS, particularly in some of the lower abundance components. Specifically, significant increases in the relative abundance of Collagen IV and VI peptides and decreased Collagen XIV were found in CDH+ vs. healthy hearts (*Figure 2, Supplementary Table 1*). High abundance proteins such as Collagen I, Fibrillin-1, and Fibronectin were not significantly different.

**Figure 1.**
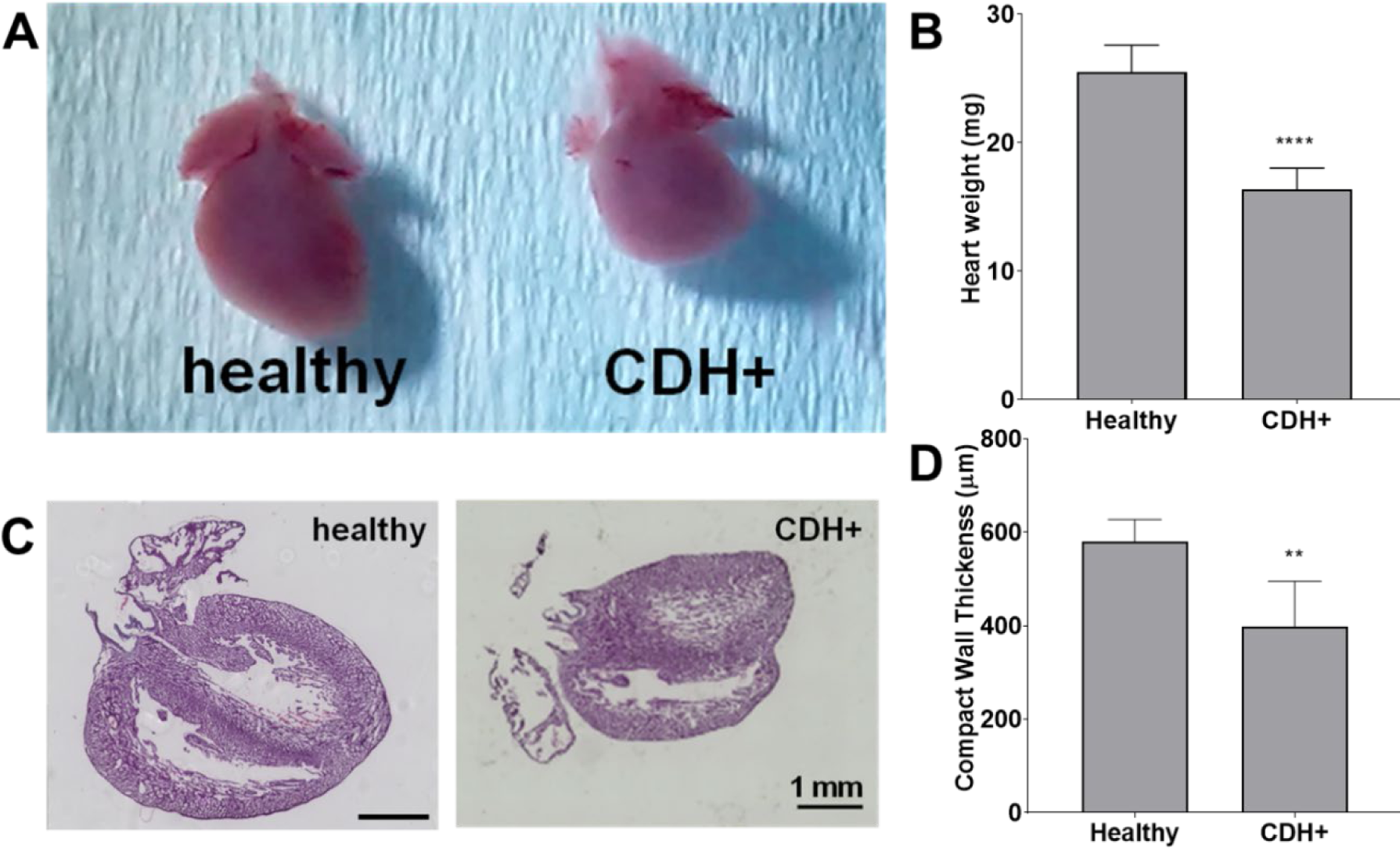
Altered heart morphology in the rat model of CDH. (A) Representative images of E21 hearts from healthy controls and nitrofen-treated CDH+ fetuses. (B) Heart mass measurements. (C) Representative H & E stained sections of healthy and CDH+ hearts. Scale bars = 1mm. (D) Measurements of left ventricle (LV) free wall thickness (n = 5).

**Figure 2.**
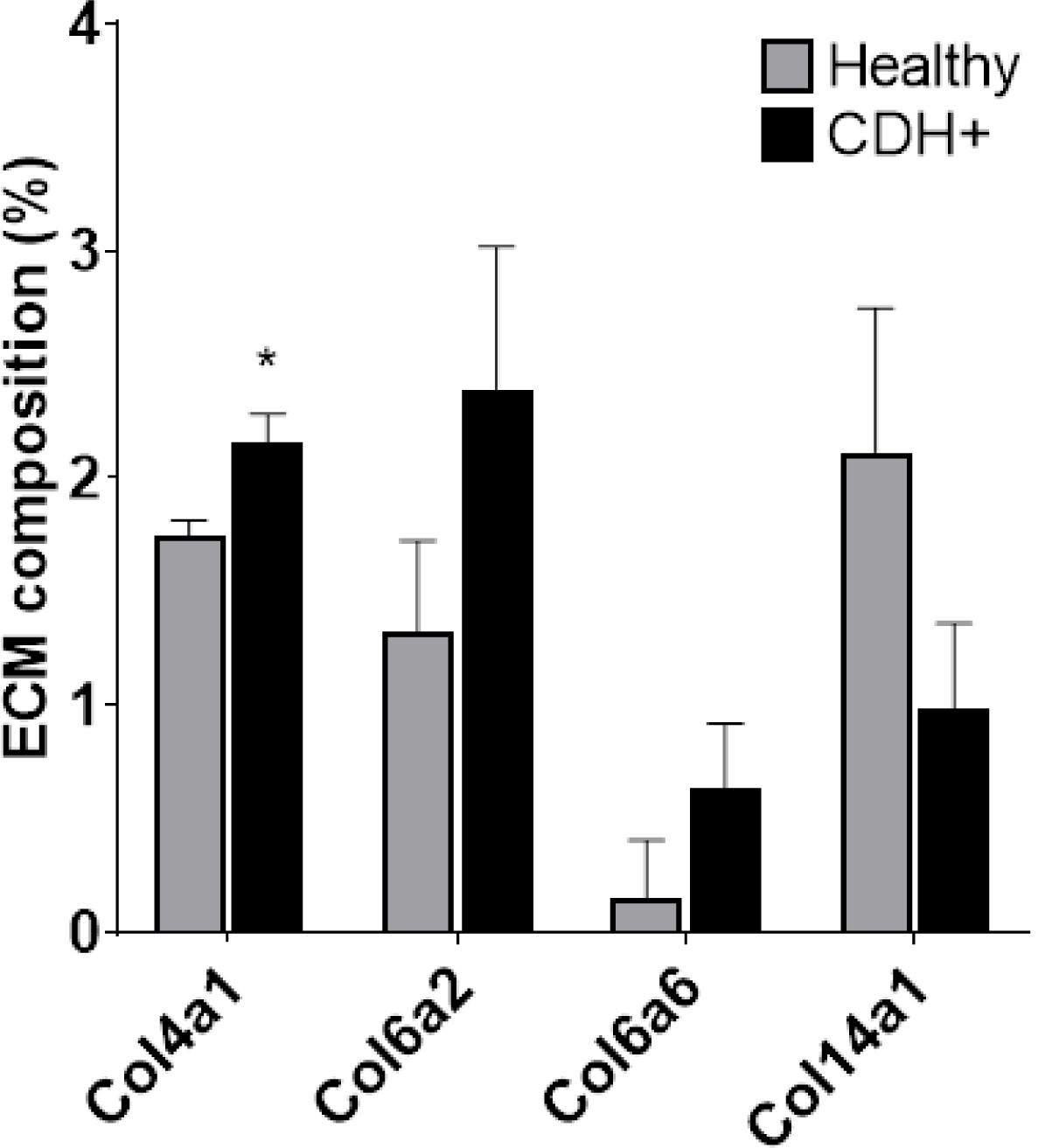
ECM alterations in CDH. Significant differences were found in low abundance proteins.

### 3.2. CDH+ hearts exhibit reduced cell proliferation and cardiomyocyte maturity

To determine whether CDH+ hearts exhibited reduced proliferation, native heart sections were stained for Ki67 to identify proliferating cells and sarcomeric α-actinin to label cardiomyocytes (*Figure 3A*). Cell proliferation was significantly decreased in CDH+ hearts compared to healthy controls (*Figure 3B*). Particularly in CDH+ hearts, many non-myocytes appeared to be Ki67+ (*Figure 3A, yellow arrows*). Therefore we determined whether proliferation was decreased specifically in the cardiomyocyte population. Although cardiomyocyte proliferation (as determined by Ki67+ α-actinin+ cells) showed a decreasing trend in CDH+ hearts, it was not statistically significant (p = 0.07) (*Figure 3C*).

**Supplementary Table 1:**
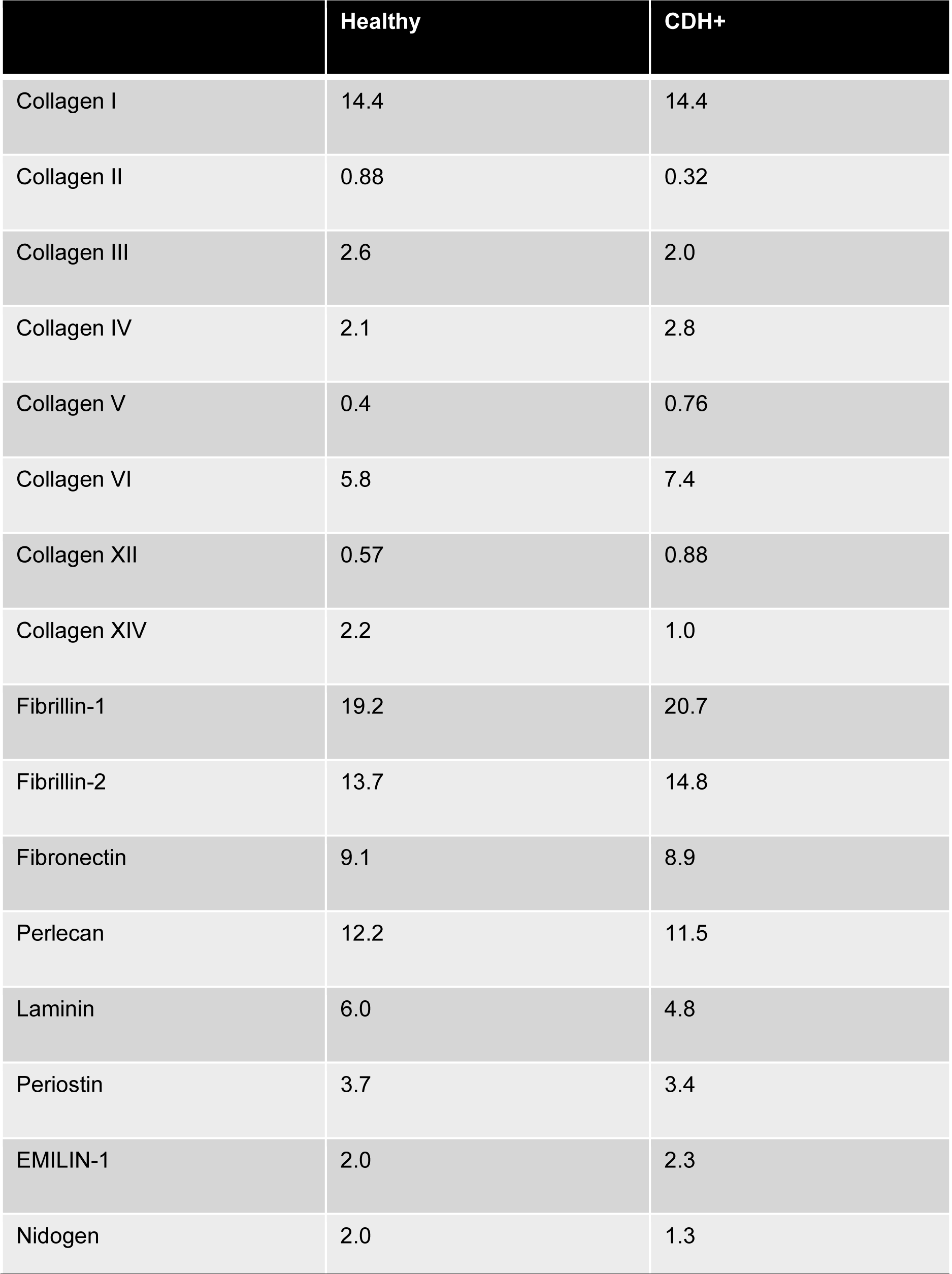
Relative ECM composition. Determined from spectral counts, data are percentages.

**Figure 3.**
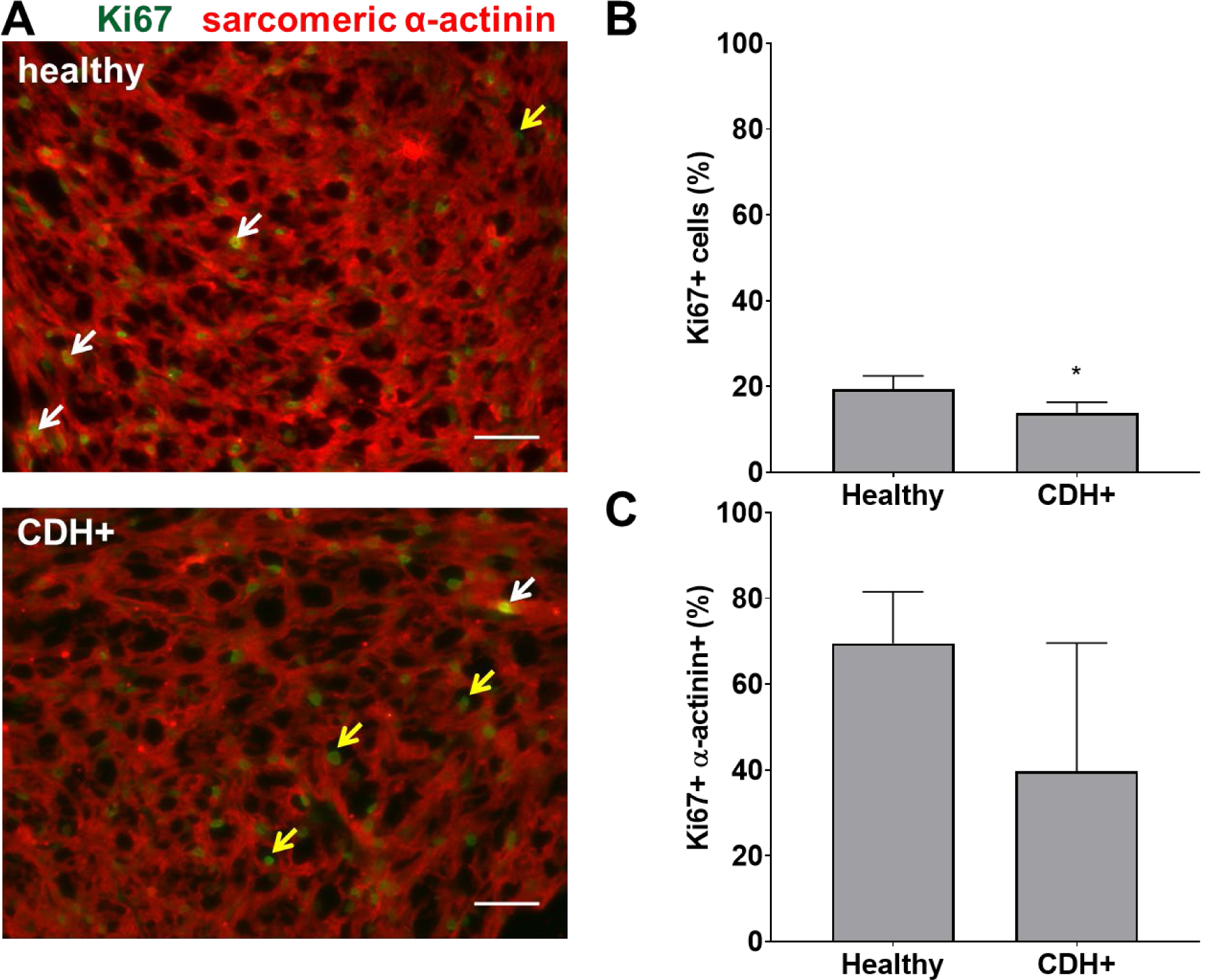
Proliferation measurements in healthy and CDH+ hearts. (A) E21 heart sections stained for Ki67 and sarcomeric α-actinin; examples of Ki67+ cardiomyocytes (white arrows), Ki67+ non-myocytes (yellow arrows), scale bars = 50 µm. (B) Total cell proliferation. (C) Cardiomyocyte-specific proliferation (the percentage of Ki67+ cells that were also α-actinin+), * = p < 0.05

To assess cardiomyocyte maturity, we analyzed sarcomere lengths and gene expression in native hearts. Average sarcomere length was significantly lower in CDH+ hearts compared to healthy (*Figure 4A, B*). In line with this data, a greater proportion of measured sarcomeres were “mature” (≥ 1.8 µm) in healthy vs. CDH+ hearts (∼80% vs. ∼50%, respectively; *Figure 4C*). Taken together, the sarcomere measurements imply that cardiomyocytes were less mature in CDH+ hearts compared to healthy. We also investigated a panel of cardiac genes and found that *mhc6*, the gene for myosin heavy chain α which is more abundant in the maturing or adult rat heart [32, 33], was significantly down-regulated in CDH+ hearts relative to healthy (*Figure 4D*). Furthermore, the ratio of mhc6 to mhc7, which has been used to assess cardiomyocyte maturity [34], was significantly decreased in CDH+ hearts (*Figure 4E*).

**Figure 4.**
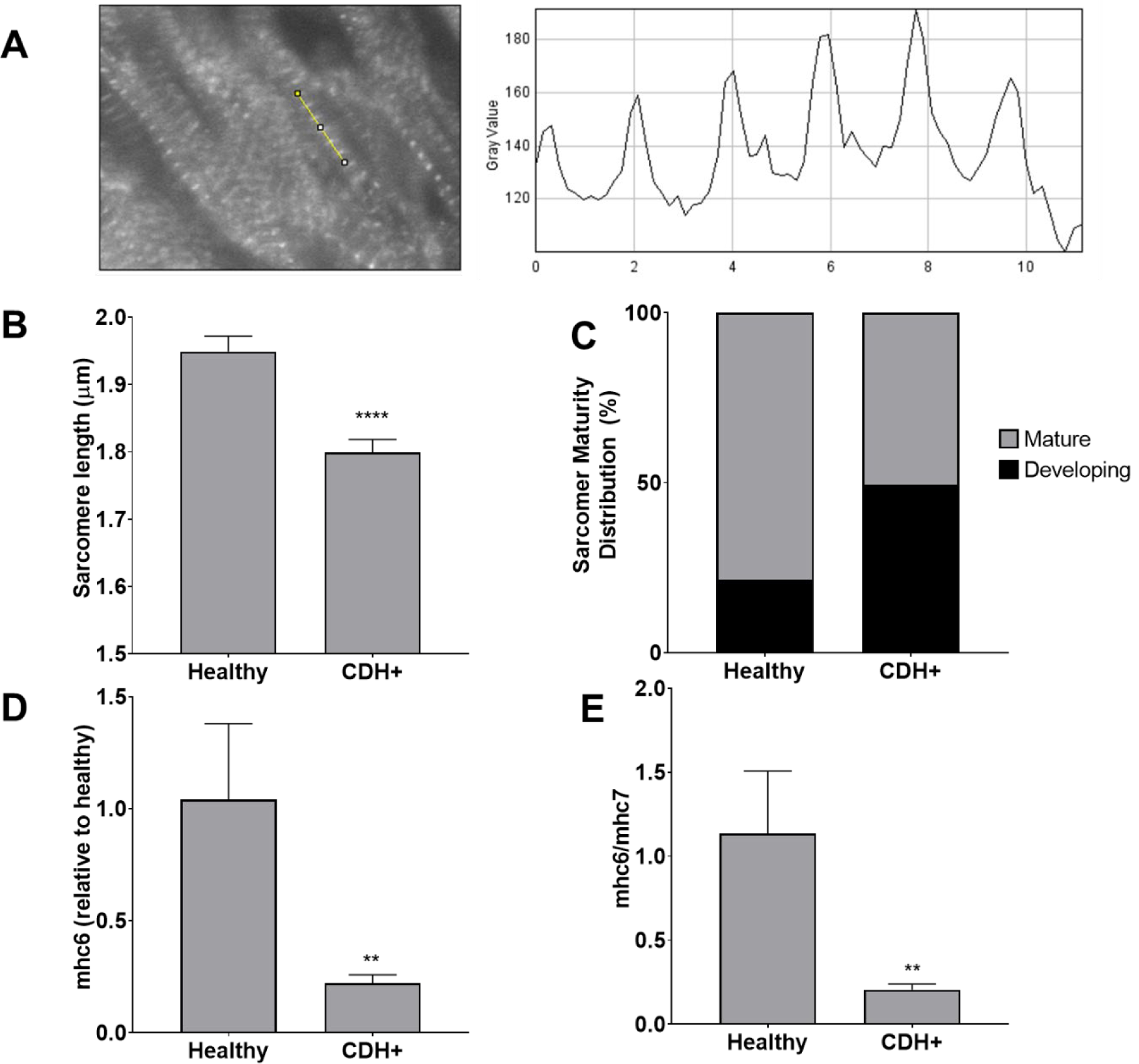
Maturation assessments in healthy and CDH+ hearts. (A) Example of sarcomere staining and measurement in native heart sections. A line is drawn perpendicular to the sarcomeres (yellow) and the profile is plotted in ImageJ. Peak-to-peak distances represent sarcomere lengths. (B) Average sarcomere length measurements (mean ± S.E.M. n > 80 individual cells per condition). (C) A greater percentage of sarcomeres were mature (≥ 1.8 µm) in healthy vs. CDH+ hearts. (D) Gene expression for αMHC (mhc6) (E) as well as the ratio of α to β isoforms (mhc6/mhc7) was decreased in CDH+ hearts (n = 4).

### 3.3. CDH+ cardiomyocytes in culture remain immature and become proliferative

To determine whether proliferation remained low or could be recovered upon removal from the hypoplastic environment, we isolated cells from CDH+ and healthy hearts and cultured on standard tissue culture plastic. After 1 day cardiomyocyte density was similar for both conditions, but after 6 days there were significantly more cardiomyocytes in the CDH+ population compared to the healthy population (*Figure 5A*). Whereas the healthy cardiomyocytes did not significantly increase in number from 1 to 6 days (fold change of 1.45), the CDH+ cardiomyocytes increased 2.4-fold (p = 0.0025). At 2 days, Ki67 was detected in 24% of the CDH+ cardiomyocytes, compared to only 12% in the healthy, demonstrating that the CDH+ cells were indeed more proliferative (*Figure 5B*). We also found that sarcomeres were significantly smaller in the CDH+ cardiomyocytes after 6 days, suggesting that these cells remained less mature than healthy cells (*Figure 5C*).

**Figure 5.**
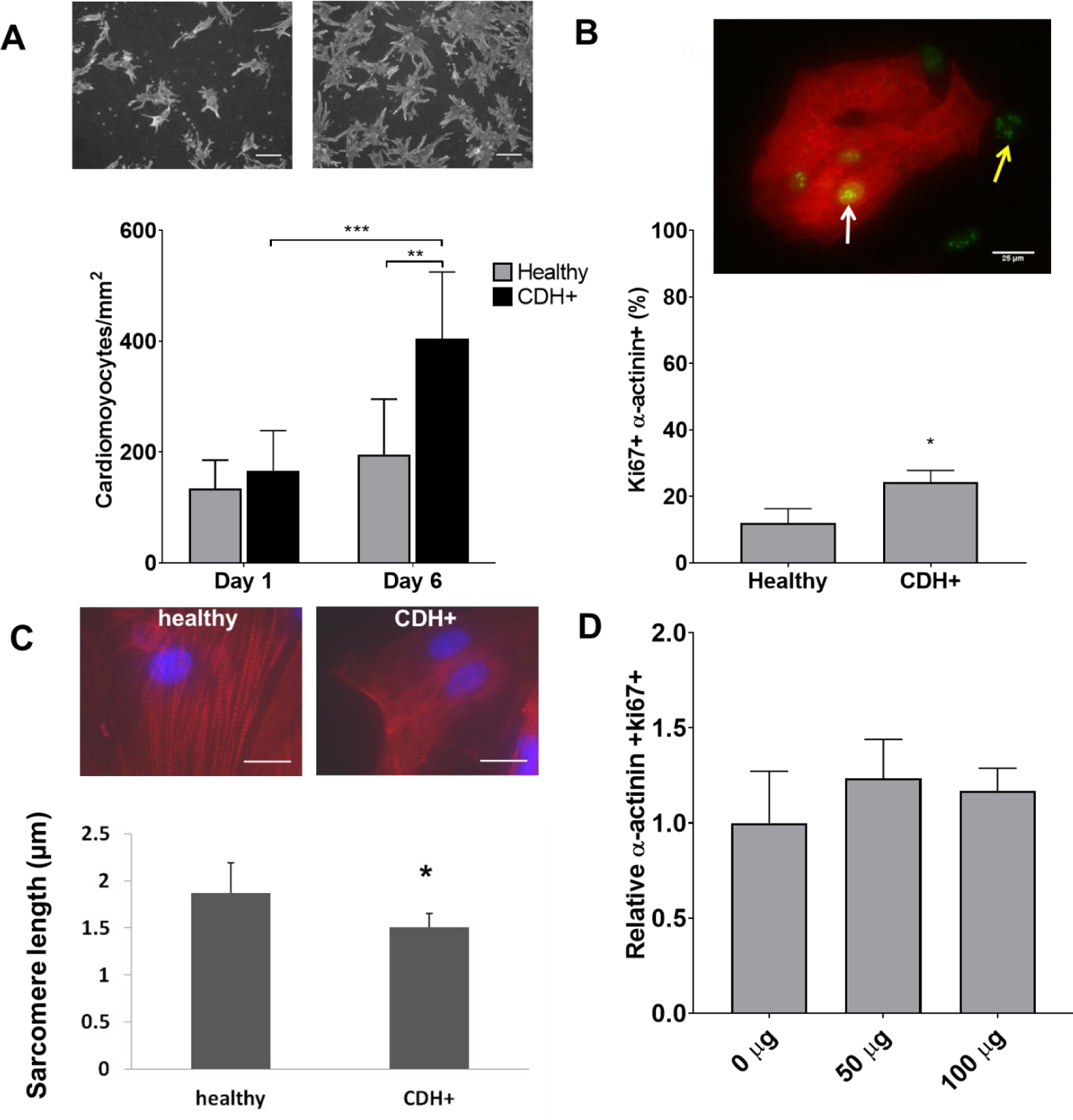
Characterization of CDH+ cardiomyocytes in culture. (A) Representative images of cardiomyocytes (α-actinin stain) after 6 days in culture (top). Cardiomyocyte density at 1 and 6 days (bottom). Graph shows mean ± SD (n = 6). (B) Cardiomyocyte proliferation (Ki67+ α-actinin+) at 2 days. Inset shows example of Ki67+ cardiomyocyte (white arrow) and non-myocyte (yellow arrow). Graph shows mean ± SD (n = 3). (C) Representative images of sarcomere staining in healthy and CDH+ cardiomyocytes and sarcomere length measurements. (D) Proliferative cardiomyocytes at 72 hours post-treatment with exogenous nitrofen. Data is normalized by relative expression of α-actinin+ and Ki67+ at 24 hours pre-treatment. Graph shows mean ± SD (n = 5).

### 3.4. Culture with exogenous nitrofen does not affect proliferation in healthy cardiomyocytes

In order to determine whether alterations in cardiomyocyte proliferation in the CDH+ animals was due to an direct effect of nitrofen, we treated cardiomyocytes with nitrofen and assessed proliferation over the course of 3 days. For these studies, freshly isolated cardiomyocytes were cultured with 0 mg, 50 mg, and 100 mg of nitrofen. After 72 hours of culture, we observed no significant differences in in ki-67+ cardiomyocytes that were treated with exogenous nitrofen compared to cardiomyocytes cultured without nitrofen (*Figure 5D)*.

### 3.5. CDH+ cardiac ECM inhibits cardiomyocyte proliferation

Given that ECM composition was altered in CDH+ hearts (*Figure 2*), we hypothesized that the ECM plays a role in the decreased proliferation observed in CDH-associated HH. Healthy and CDH+ cardiomyocytes were seeded onto healthy and CDH+ heart-derived ECM and cultured with serum-free medium. After 4 days, cardiomyocyte numbers were significantly higher on healthy ECM compared to CDH+ ECM for both healthy and CDH+ cardiomyocytes (*Figure 6A, B*). CDH+ cardiomyocytes were generally more proliferative than their healthy counterparts on either ECM. However, culture on CDH+ ECM resulted in significantly decreased cardiomyocyte proliferation for both healthy and CDH+ populations (*Figure 6C*). This data suggests that changes in ECM composition present in CDH+ hearts inhibits cardiomyocyte proliferation in CDH-associated HH.

**Figure 6.**
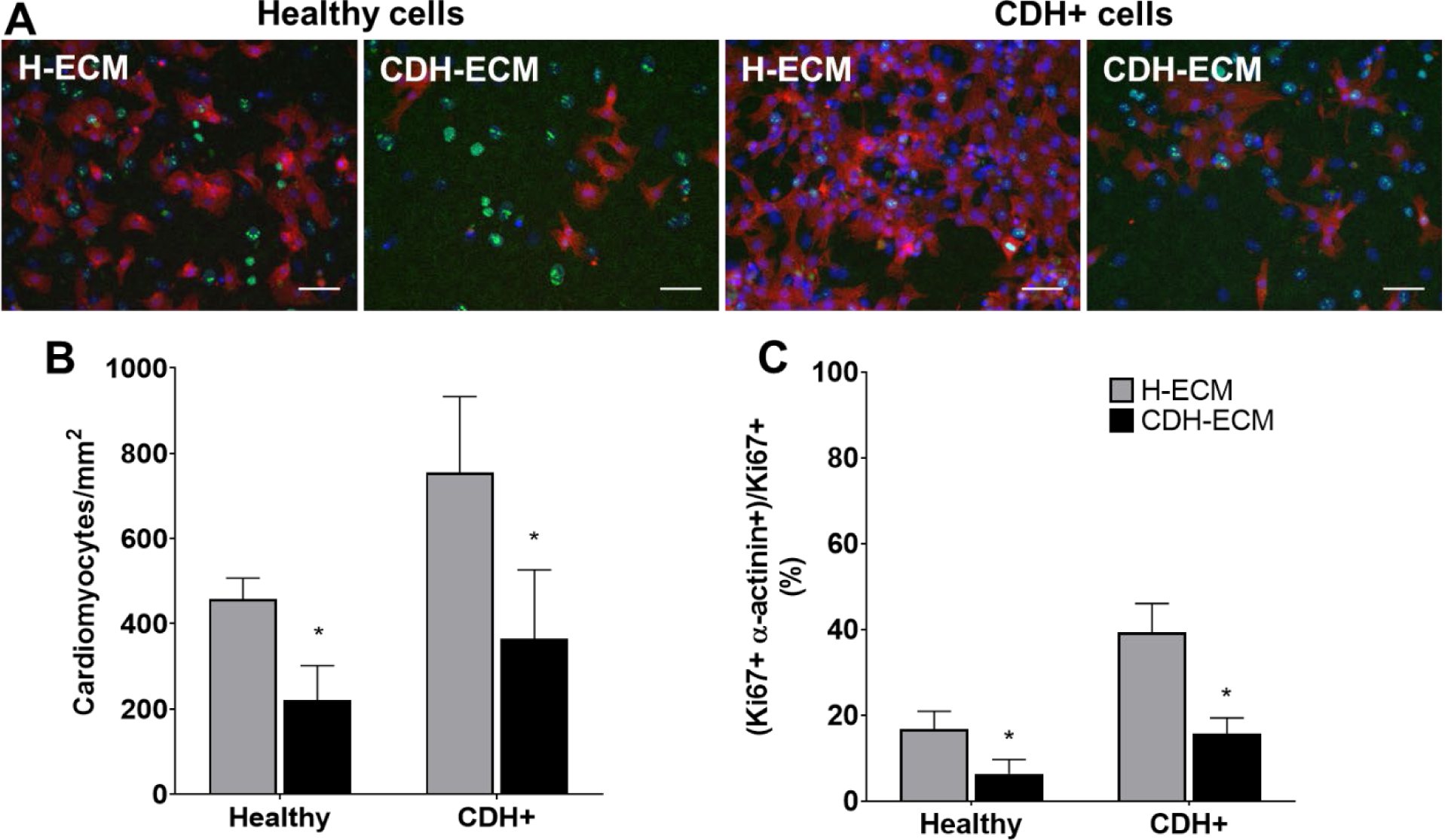
Cardiomyocyte response to ECM. (A) Representative images of cells on ECM derived from healthy hearts (H-ECM) and CDH+ hearts (CDH-ECM), stained for Hoechst (blue), Ki67 (green), and cardiac α-actinin (red). (B) Cardiomyocyte density and (C) cardiomyocyte proliferation were significantly decreased in both healthy and CDH+ populations on CDH-ECM at 4 days. Graphs show mean ± SD (n = 3).

### 3.6. Cyclic mechanical loading promotes maturation of CDH+ cardiomyocytes

Compression of the heart likely impedes the ability of the heart to undergo normal stretch in CDH [19]. We hypothesized that mimicking physiological stretch *ex vivo* could promote CDH+ cardiomyocyte proliferation and maturation. For these studies, we used a frequency of 1 Hz at 5% amplitude. We did not find a significant effect of stretch on proliferation at the time point studied (3 days of stretching, a total of 7 days in culture). However, CDH+ cardiomyocytes had significantly longer sarcomeres after stretching compared to static controls. In contrast, sarcomere lengths in healthy cells were unaffected by stretch (Figure 7). These results show that cyclic mechanical stretch promoted sarcomere maturation in CDH+ cardiomyocytes.

**Figure 7.**
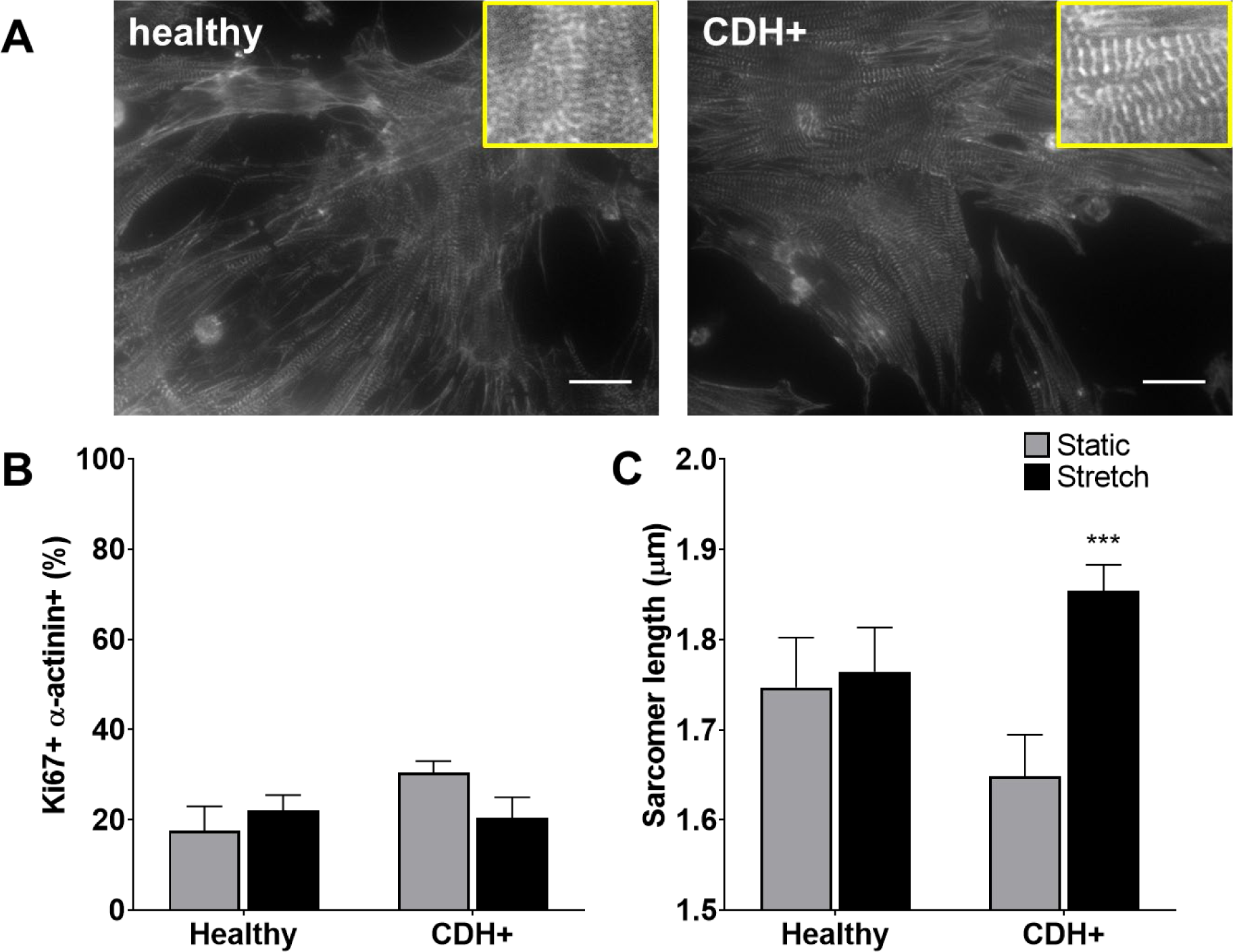
Cardiomyocyte responses to stretch. (A) Representative images of stretched cardiomyocytes (α-actinin stain). Insets show sarcomeres. Scale bars = 25 µm. (B) Cardiomyocyte proliferation was not significantly affected by stretch. Graph shows mean ± SD (n ≥ 2). (C) CDH+ cardiomyocytes underwent lengthening of sarcomeres in response to stretch. Graph shows mean ± SEM (n ≥ 23);

## 4. Discussion

The nitrofen model of CDH in rats is a valuable tool for studying molecular and cellular alterations in HH which are difficult in humans. Although a few groups have looked at changes in growth factors [19, 35, 36], and cardiac and ECM genes [19, 37, 38] in the nitrofen model of CDH, CDH+ cardiac cells have not been cultured or subjected to specific engineered environments to study their response to biophysical cues. The key novel findings of this study are: 1) cell proliferation and cardiomyocyte maturity were decreased in CDH+ hearts compared to healthy-importantly these effects were not caused by a direct effect of nitrofen on cardiomyocytes in the developing hearts; 2) in culture, CDH+ cardiomyocytes remained immature with increased proliferative potential compared to healthy cardiomyocytes; 3) ECM derived from CDH+ hearts significantly reduced both healthy and CDH+ cardiomyocyte proliferation compared healthy cardiac ECM; and 4) cyclic mechanical stretch promoted sarcomere maturation in CDH+ cardiomyocytes.

Given that the healthy heart undergoes a transition from hyperplastic to hypertrophic growth soon after birth [39, 40], cardiomyocyte proliferation and maturation are usually considered to be inversely correlated. However, we found that CDH+ hearts had smaller sarcomeres and reduced *mhc6* expression, suggesting a less mature state, while also having decreased proliferation compared to healthy controls. Numerous external factors can influence cell behavior, such as growth factor signaling, the ECM, tissue stiffness, and the dynamic mechanical environment [6-10, 12, 41]. While some of these cues are known to be altered in HH, it is unclear how their complex interactions could lead to concurrent decreased proliferation and maturity in the CDH+ heart. It was initially surprising to find that in culture the proliferation of CDH+ cardiomyocytes significantly exceeded that of healthy cardiomyocytes. It appears that the immature state of CDH+ cardiomyocytes is advantageous for recovered growth, as the removal of the hypoplastic environment allowed the cells to undergo a proliferative “burst” that would not be achievable by more mature myocytes. A similar mechanism may exist in young patients who undergo CDH repair, as recovery of heart dimensions have been observed in patients with mild to moderate HH [3]. Given recent studies of young human hearts [8, 42], therapeutic strategies which promote cardiomyocyte proliferation will likely be most effective during early life.

Cardiac ECM influences cardiomyocyte proliferation [6, 7] but has not been well studied in the context of HH disorders. We found that culture on CDH+ ECM led to reduced proliferation of both healthy and CDH+ cardiomyocytes. This was intriguing since there were significant changes in only a small fraction of ECM components (Collagens IV, VI, and XIV, which were each < 3% of the ECM) relative to total composition. However, these proteins have important roles that could affect how cells sense and respond to the ECM [43-47]. In CDH-associated HH and HLHS, two studies suggest that the ECM in hypoplastic hearts is less mature compared to healthy hearts [19, 20, 47]. Specifically, Guarino et al found reduced procollagen and tropoelastin in CDH+ fetal rat hearts and Davies et al observed increased fibronectin and decreased collagen in HLHS hearts compared to non-HLHS patients. Tao et al. found that Col14a null mice displayed defects in ventricular morphogenesis and had significantly increased cardiomyocyte proliferation postnatally but a similar trend as in our CDH model towards decreased cardiomyocyte proliferation at E11.5 [46]. Similarly, a recent study by Bousalis et al. observed increased expression of fibronectin, collagen IV, and integrin β-1 within *Nkx2-5* mutant embryonic mouse hearts compared to wild type hearts [47]. Interactions between fibronectin and integrins can lead to downstream signaling cascades that control cell spreading, migration, and proliferation [48, 49]. Our findings are counter-intuitive, as immature ECM has generally been found to promote proliferation [6-8, 50]. The role of the ECM is more complex; future studies should explore the roles of specific peptides or interactions among ECM components to elucidate the nuances of the ECM in HH pathology and cardiomyocyte proliferation.

Normal stretching of the heart wall is diminished in CDH due to compression by visceral organs [19]. We found that CDH+ cardiomyocytes had lengthened sarcomeres when stretched using a custom flexible membrane apparatus compared to static conditions; healthy cells, which were already relatively mature, did not further lengthen their sarcomeres under these cyclic loading conditions. Cyclic stretch has been used as a strategy to drive cardiomyocyte maturation in a number of tissue engineering applications [51, 52]. Diminished mechanical movement experienced by cardiomyocytes within CDH+ hearts could be a potential mechanism of arrested cardiomyocyte maturity in CDH-associated HH and has also been implicated in HLHS due to reduced blood flow in the left side of the heart [53]. The role of mechanical forces is intriguing when contrasting CDH-associated HH and HLHS. Although these two HH defects share some similar features such as reduced growth factors, altered ECM, and decreased cardiac transcription factors [5, 19-21, 54, 55], the outcomes after surgical repair are vastly different. Repair of CDH leads to removal of the compressive forces on the heart by intruding organs, and thus cardiac growth often normalizes [3]. This is not the case for HLHS: restoration of blood flow via fetal aortic valvuloplasty does not lead to recovered left ventricular growth [15]. A deeper understanding of the specific underlying pathological mechanisms of these different forms of heart hypoplasia is needed.

In conclusion, we have found that cardiomyocytes from CDH+ hearts are capable of robust proliferation, likely a result of their immature phenotype maintained during heart development. Our studies also point to altered ECM and mechanical forces as important environmental regulators of cardiomyocyte state in heart hypoplasia. Future work aimed at understanding these mechanisms could lead to novel biomechanical signaling-based therapies to improve cardiac growth and function in children born with hypoplastic hearts.

## Acknowledgments

The authors thank Professor John Asara and the Mass Spectrometry Core Laboratory (Beth Israel Deaconess Medical Center) for running LC-MS/MS on ECM samples.

